## Supplementary Materials for "The refinement paradox and cumulative cultural evolution: collective improvement in knowledge favors conformity, blind copying and hyper-credulity"

**This PDF includes:**

Figs. S1-S8
Tables S1-S5

### Supplementary Figures

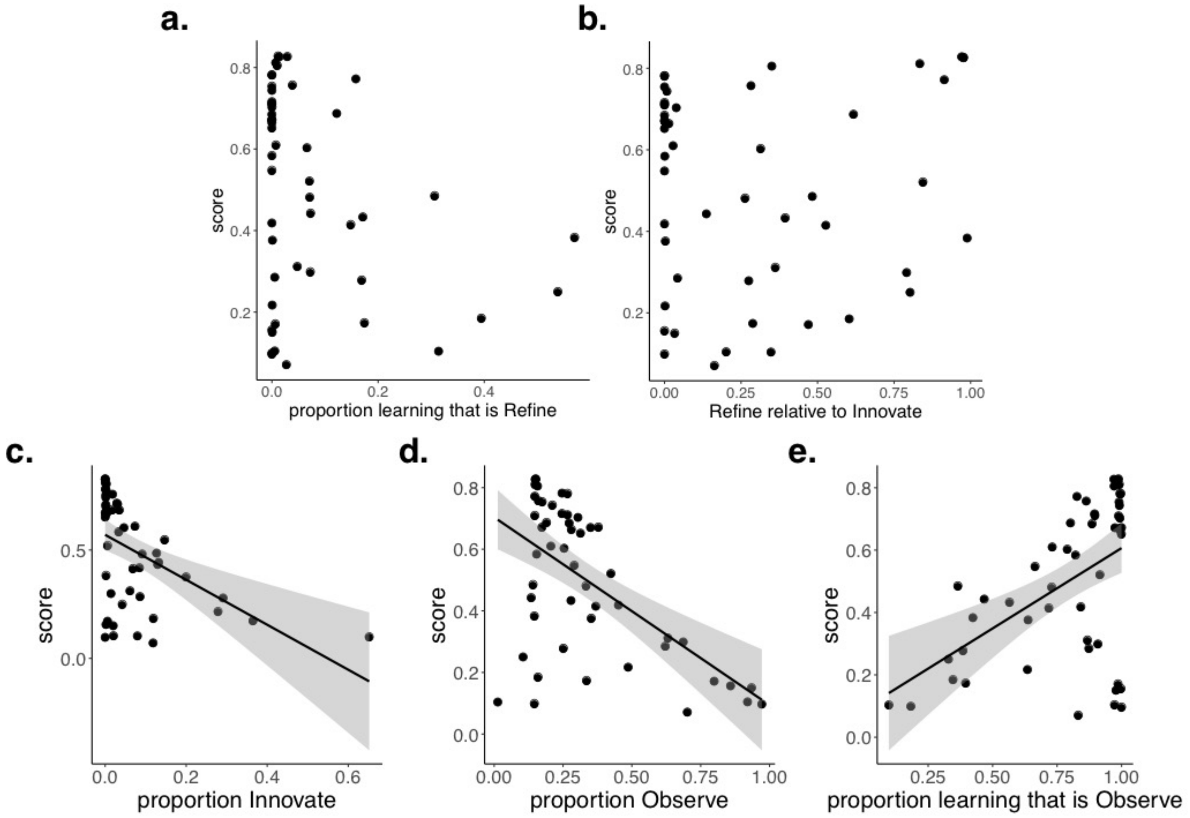


**Fig. S1.** Relationship between score and (**a**) amount of REFINE as a proportion of all learning moves; (**b**) proportion of just REFINE and INNOVATE learning moves; (**c**) proportion of INNOVATE moves; (**d**) proportion of OBSERVE moves and (**e**) proportion of learning moves that are OBSERVE (and not INNOVATE or REFINE).

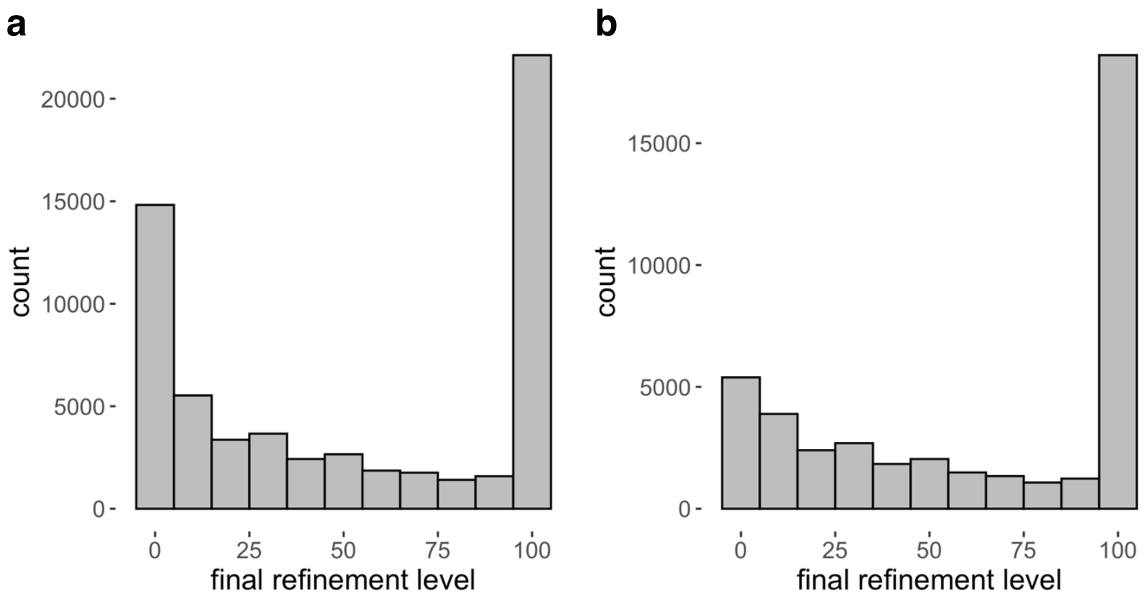


**Fig. S2.** Distribution of final maximum refinement level in the tournament (**a**) in all simulations, and (**b**) in simulations that include only entries that use REFINE. Refinement levels above the value of 8 typically lead to payoffs that exceed the maximum basic payoff (Fig. 1). The maximum refinement level achieved across simulations varied substantially and showed a bimodal distribution.

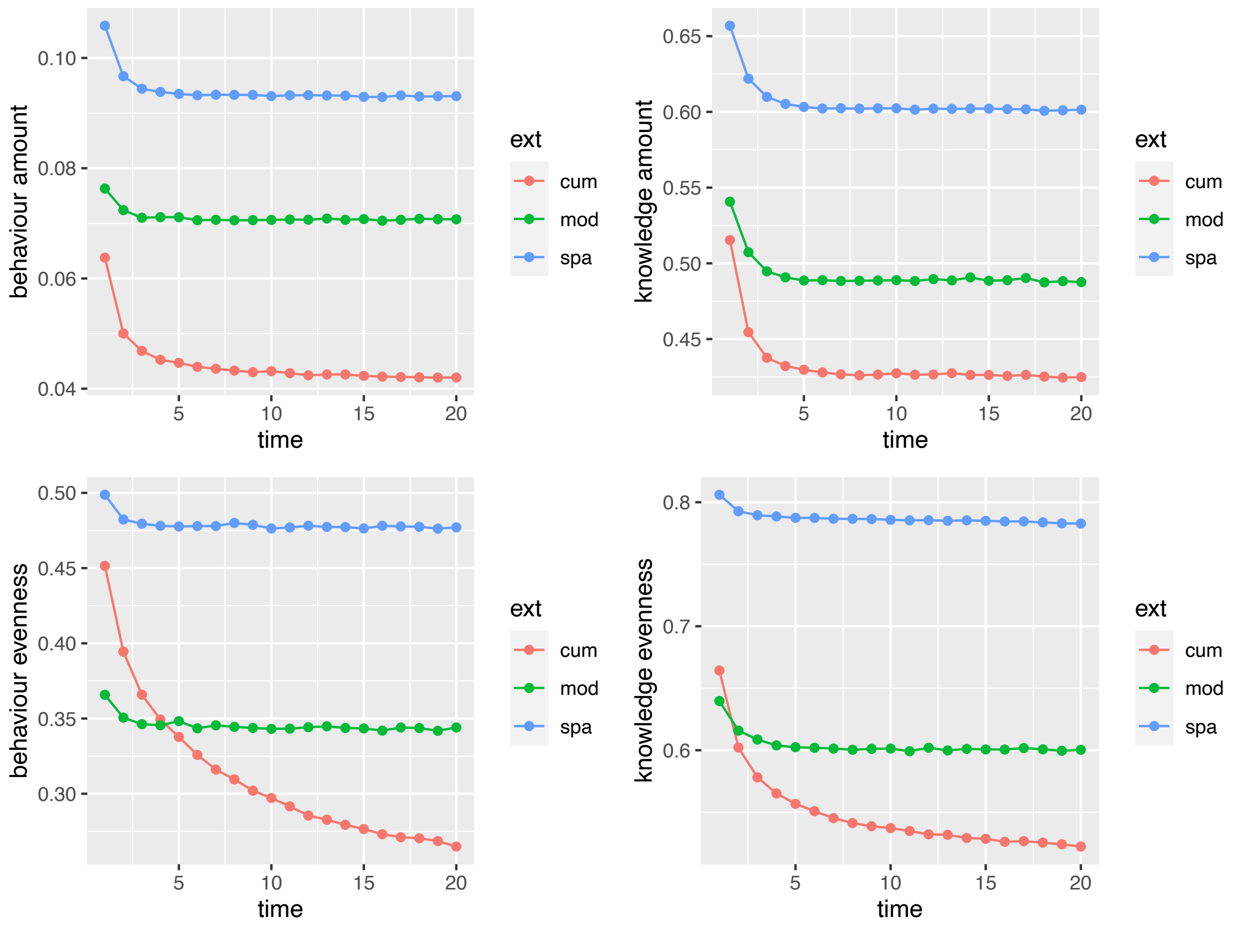


**Fig. S3.** Cultural diversity measures across extensions for Stage I. Cumulative culture leads to decreasing diversity in both the behaviors performed and behaviors known about, as populations converge on a small number of heavily refined, high-payoff behaviors.

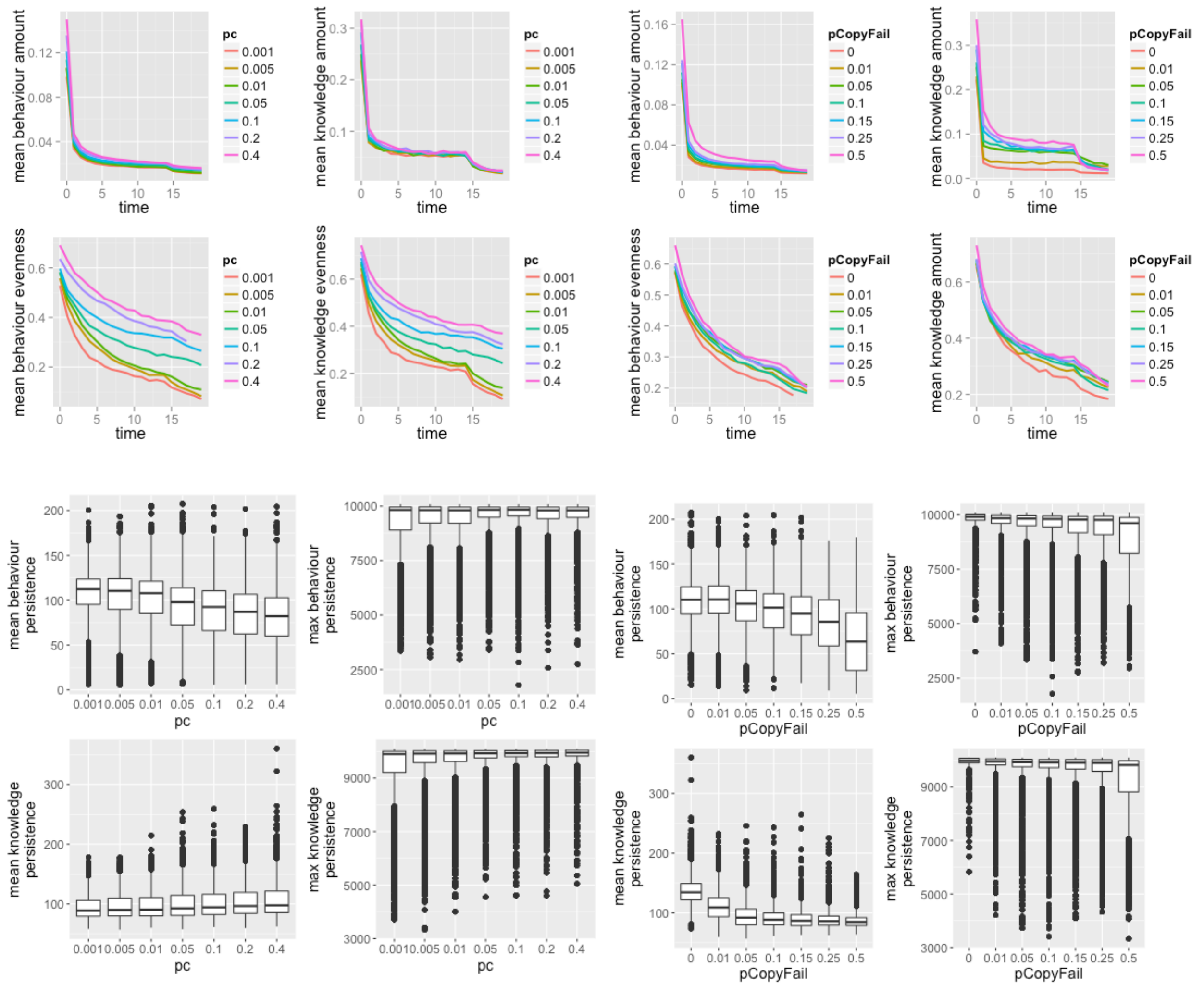

**Fig. S4.** Cultural diversity measures as a function of $p_c$ and $p_{copyFail}$ in Stage 2. Timelines of amount and evenness are presented as line charts, while the boxplots illustrate mean values for mean and maximum persistence.

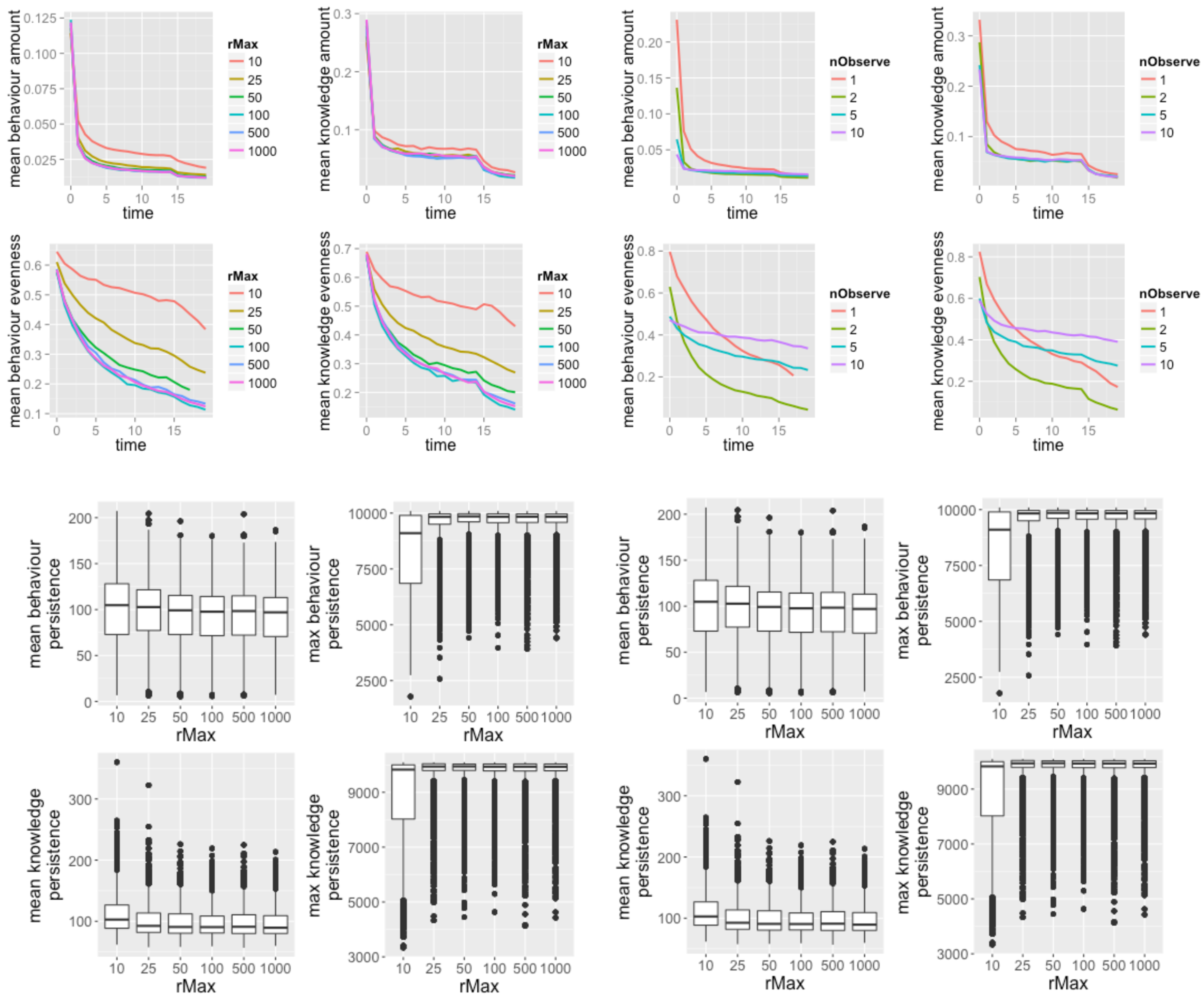


**Fig. S5.** Cultural diversity measures as a function of $r_{max}$ and $n_{Observe}$, in Stage 2.

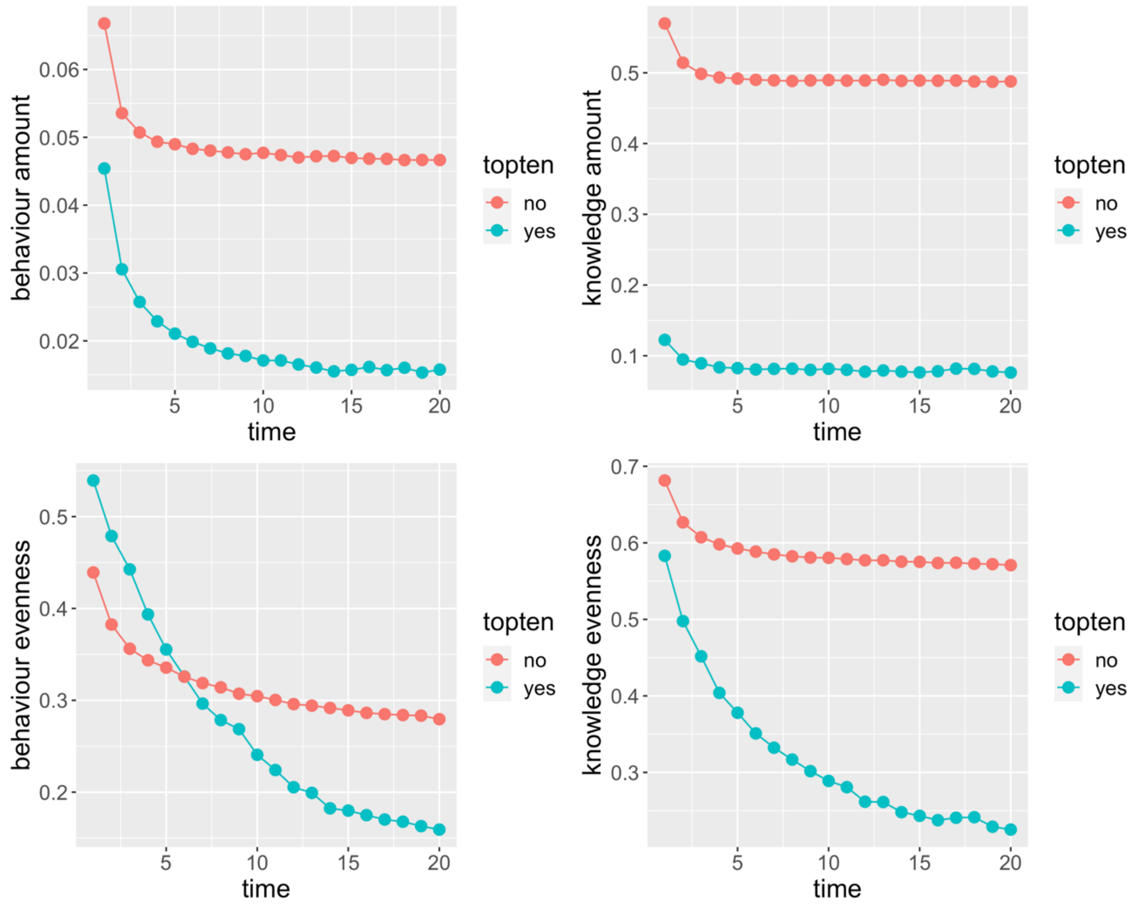


**Fig. S6.** Amount and evenness of both behavior and knowledge in simulations that only include top ten entries (teal) and simulations with the rest of the 41 entries (red), in Stage I (cumulative extension).

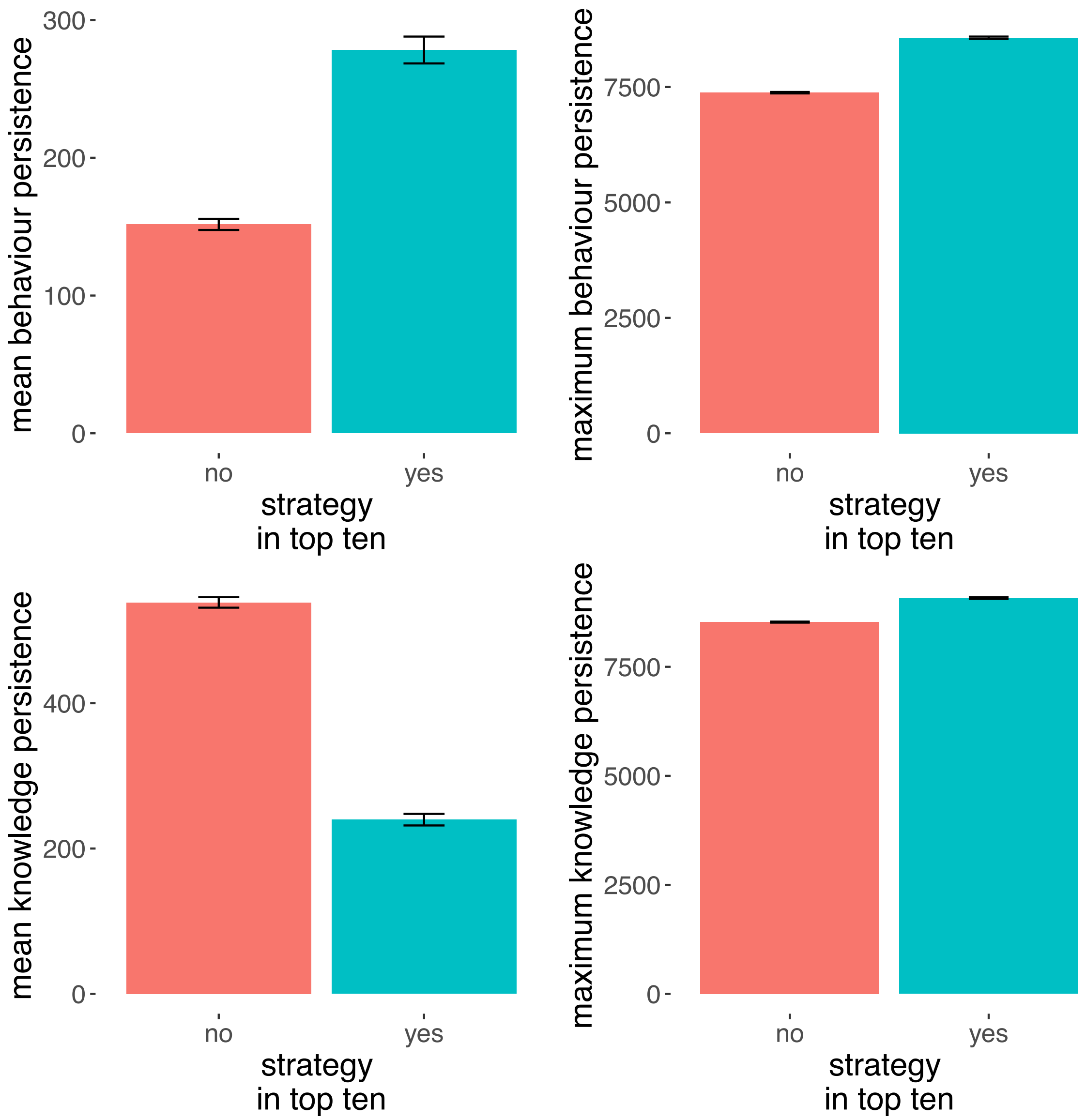


**Fig. S7.** Mean and maximum persistence of both behavior and knowledge, in simulations that only include the top ten entries (teal) and simulations with the rest of the 41 entries (red), in Stage I (cumulative extension).

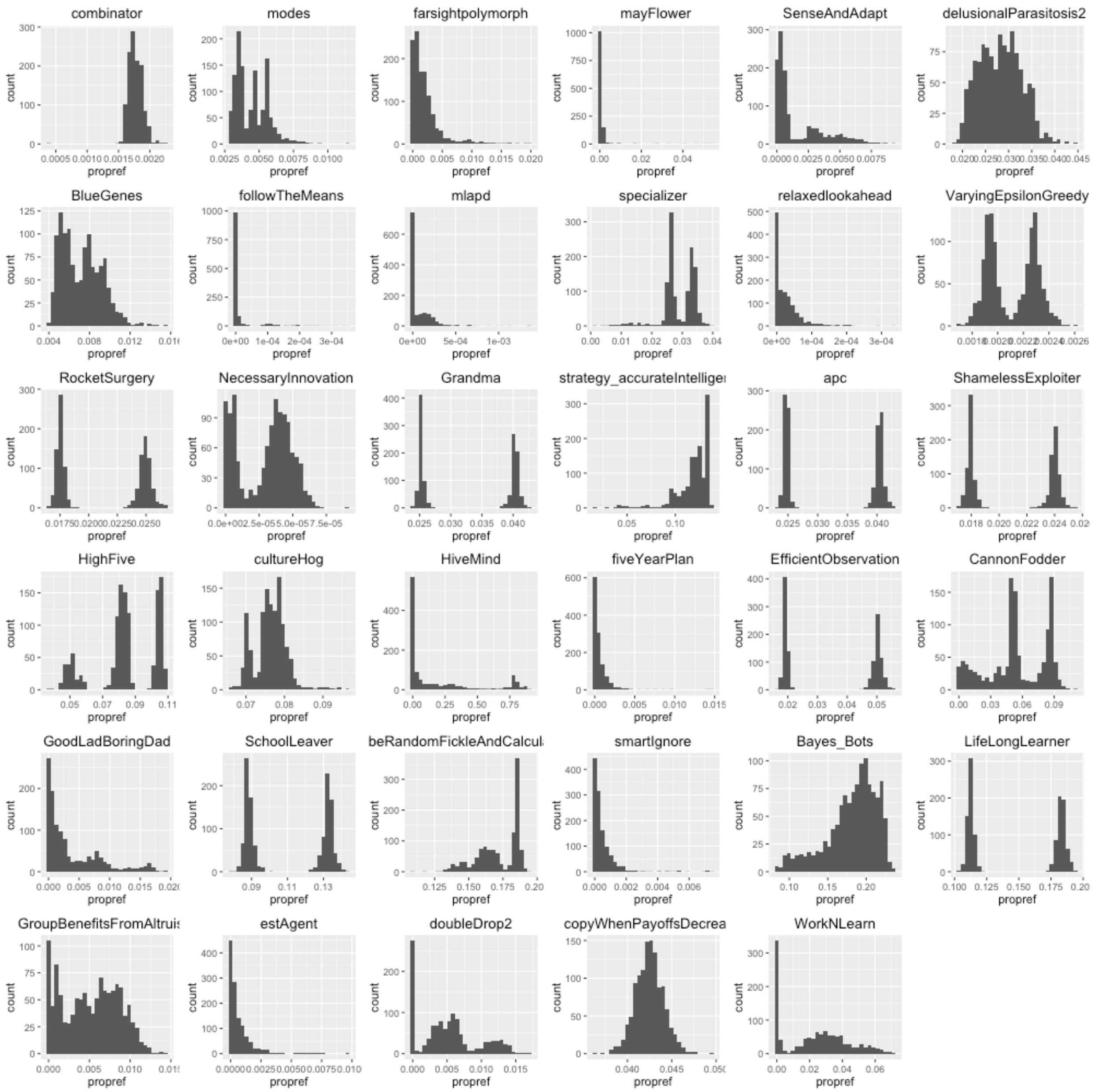


**Fig. S8.** Distribution of mean proportion REFINE moves per simulation for each entry, ordered from the top left by score in Stage 1, in descending order. This plot only includes the 35 entries that used the REFINE move. Please note the difference in scale on the x axis. The top four entries were also the top performers in Stage 3, which establishes that the best-performing entry used REFINE at low levels.

### Supplementary Tables

**Table S1.** Model averaged parameter estimates and summed Akaike weights (across an all-subsets model set of linear regression models with Gaussian error) for Stage 1 cumulative extension. Other extensions give similar findings (see Table S2). The $AIC_c$ best model (adjusted R-squared=0.86) contained *pLearnRefine* and *pObserve*, as well as *meanPayDiffObserve*, *meanBetweenLearn* and *doRefine*. The first two predictors were included in all of the 13 well-supported ($\Delta AIC_c$<2) models; while the others were in 9, 6 and 7 respectively.

| Term | Definition | Estimate | s.e | $\sum\omega_i$ |
|---|---|---|---|---|
| pLearnRefine | Proportion of all moves that were either REFINE, INNOVATE, or OBSERVE i.e. not EXPLOIT. | -0.798 | 0.078 | 1.000 |
| pObserve | Proportion of learning moves (OBSERVE +INNOVATE) that were OBSERVE. | 0.379 | 0.077 | 1.000 |
| doRefine | Does the entry play, or have the possibility of playing the REFINE move? (scored 0/1) | -0.057 | 0.035 | 0.529 |
| meanPayDiffObserve | The average difference in payoffs between EXPLOIT moves immediately preceding an OBSERVE move and the next EXPLOIT move after it. | -0.001 | 0.001 | 0.521 |
| meanBetweenLearn | The average number of rounds between learning moves. | -0.004 | 0.003 | 0.406 |
| pFailEst | Does the entry estimate $p_{copyFail}$, the probability that observing fails? (scored 0/1). | -0.070 | 0.058 | 0.354 |
| checkPayDist | The entry uses some aspect of the agent's historical payoffs in deciding how to move – either the maximum or mean of all the payoffs in the history, or possibly some subset of the entire history. | -0.036 | 0.035 | 0.309 |
| payDrop | Does the entry have a specific rule for dealing with the situation in which the payoff for a known behaviour drops below some threshold after being used in an EXPLOIT move? (scored 0/1) | -0.030 | 0.032 | 0.295 |
| pcEst | Does the entry estimate $p_c$, the probability of environmental change? (scored 0/1) | 0.032 | 0.036 | 0.276 |
| lineLength | The number of lines in the Python source code for the entry. | <0.001 | <0.001 | 0.257 |
| checkRefInc | Does the entry calculate the payoff increments before and after REFINE has been played using a particular behaviour? (scored 0/1) | 0.029 | 0.040 | 0.249 |
| discountsOld | Does the entry perform some sort of discounting of historical information, either by truncating histories or weighting older data using some function of the age of the information? (scored 0/1) | 0.020 | 0.041 | 0.226 |
| nObsEst | Does the entry estimate the number of other agents surveyed in an OBSERVE move, the simulation parameter $n_{observe}$? (scored 0/1) | 0.010 | 0.036 | 0.206 |
| poly | An entry is described as polymorphic if the agent deploying it may be of more than one "type" (e.g., sometimes producer and sometimes scrounger). For instance, some entries feature a random choice made at birth and signalled in the history by a particular move, which then defines their behaviour. In this way, a population of agents may be split into different groups, each playing different entries. | 0.007 | 0.041 | 0.200 |
| meanRoundsToExploit | The average number of rounds between the birth of an agent with this entry and that agent's first EXPLOIT move | 0.001 | 0.003 | 0.199 |

**Table S2.** Results from a linear mixed model predicting score as a function of the interaction between whether an environment was refined or not and whether and entry used refine or not, with a varying intercept for entry identity, using data for all 51 entries in the tournament. The model definition: score ~ environment type*entry type + 1|entry, assuming a non-refine entry in a non-refined environment as baseline.

| Variable | β coefficient | Standard error | t-value |
|---|---|---|---|
| **Intercept** | **0.568** | **0.064** | **8.851** |
| **refined environment** | **0.010** | **0.006** | **1.616** |
| **refine entry** | **-0.175** | **0.077** | **-2.261** |
| **refined environment * refine entry** | **0.161** | **0.007** | **22.115** |

**Table S3.** Results from a similar linear mixed model with a varying intercept for entry identity, using data for 20 entries, the top-scoring 10 entries that used REFINE and the top-scoring 10 entries that did not use REFINE. Model definition: score ~ environment type*entry type + 1|entry, assuming a non-refined entry in a non-refined environment as baseline.

| Variable | β coefficient | Standard error | t-value |
|---|---|---|---|
| **Intercept** | **0.706** | **0.017** | **40.891** |
| **refined environment** | **0.068** | **0.007** | **8.913** |
| **refine entry** | **0.044** | **0.024** | **1.826** |
| **refined environment * refine entry** | **0.029** | **0.020** | **2.890** |

**Table S4.** Definitions of new terms

| Term | Definition |
|---|---|
| Learning move | Any move used to acquire new behavior or revise present behavior, i.e. all moves but EXPLOIT |
| Refine entry | Entries that make use of the REFINE move |
| Refined environment | Environments in which the refinement level of at least one behavior has reached the maximum level, 100 |
| 'Clever' refiner | Comprised the 6 entries that were placed in the top 10, consistently played REFINE, and used REFINE in a strategic manner. |

**Table S5.** Simulation parameters

*Stage I*
*Single extension pairwise*

| Parameter | Values |
|---|---|
| $p_c$ | {0.001, 0.01, 0.1} |
| $n_{observe}$ | {1, 5} |
| $r_{max}$ | 100 |
| $p_{copyFail}$ | 0.05 |

*Stage II*
*Single extension melee*

| Parameter | Values |
|---|---|
| $p_c$ | {0.001, 0.005, 0.01, 0.05, 0.1, 0.2, 0.4} |
| $n_{observe}$ | {0, 0.01, 0.05, 0.1, 0.15, 0.25, 0.5} |
| $r_{max}$ | {1, 2, 5, 10} |
| $p_{copyFail}$ | {10, 25, 50, 100, 500, 1000} |

*Stage II*
*All extension melee*

| Parameter | Values |
|---|---|
| $p_c$ | {0.001, 0.005, 0.01, 0.05, 0.1, 0.2, 0.4} |
| $n_{observe}$ | {0, 0.01, 0.05, 0.1, 0.15, 0.25, 0.5} |
| $r_{max}$ | {1, 2, 5, 10} |
| $p_{copyFail}$ | {10, 25, 50, 100, 500, 1000} |
